# Chromosome X Dosage Modulates Development of Aneuploidy in Genetically Diverse Mouse Embryonic Stem Cells

**DOI:** 10.1101/2024.06.29.601344

**Authors:** Alexander Stanton, Selcan Aydin, Daniel A. Skelly, Dylan Stavish, Kim Leonhard, Seth Taapken, Erik McIntire, Matthew Pankratz, Anne Czechanski, Tenneille Ludwig, Ted Choi, Steven P. Gygi, Ivana Barbaric, Steven C. Munger, Laura G. Reinholdt, Martin F. Pera

**Author notes:** Correspondence (M.F.P.).

## Abstract

The genetic integrity of pluripotent stem cells (PSC) is critical to their applications in research and therapy, but it is compromised by frequent development of structural chromosome variants associated with malignancy. Many cell lines exhibit remarkable genetic stability, but little is known about the basis of the known variation in genomic integrity amongst different PSC isolates. Here we identify aneuploidies using RNA-seq and proteomics data from a panel of mouse embryonic stem cell (mESC) lines derived from 170 Diversity Outbred mice. We found 62 lines with detectable aneuploid subpopulations and a subset of originally XX lines that lost one Chromosome X (XO). Strikingly, a much lower proportion of XX lines were aneuploid, compared to XY or XO lines. Two single-cell RNA-seq data sets demonstrated that aneuploid XY DO mESC also show lower Chromosome X gene expression, and a prospective study confirmed that XY mESC accumulate higher aneuploid proportions in culture than isogenic XX lines. We identify potential mechanisms for this protective effect of X chromosome dosage, including our findings that the lines with two active X Chromosomes have a higher proportion of 2-cell-like cells, a state associated with maintenance of genetic integrity of mESC, and that they show differential expression of X-linked tumor suppressor genes associated with the DNA damage response.

**Highlights:**

- First genetic analysis of predisposition to aneuploidy in pluripotent stem cell cultures
- Chromosomal regions duplicated in aneuploid mouse embryonic stem cells are syntenic with regions overrepresented in human pluripotent stem cell lines bearing recurrent genetic abnormalities
- X-Chromosome dosage strongly influences susceptibility to aneuploidy in mouse embryonic stem cells and to a lesser degree in human pluripotent stem cells
- XX mouse embryonic stem cell lines show a higher proportion of cells in 2 cell-like state and higher expression of tumor suppressor genes associated with DNA damage response

## INTRODUCTION

One of the most significant obstacles to the laboratory or clinical use of pluripotent stem cells (PSC) is their tendency to develop recurrent genetic abnormalities during prolonged culture. Structural or numerical chromosome abnormalities are the most commonly observed variants. The aneuploidies that occur can be segmental or whole chromosome copy number variations. In human PSC, it is common to see duplications of Chromosomes 1, 7, 12, 14, 17, 20, X, as well as smaller regions of these and some other chromosomes^1–8^ whereas in mice it is common to see duplications of Chromosomes 1, 6, 8, and 11 as well as loss of one sex chromosome (X in female lines and Y in male lines)^9–14^. Despite their disparate nature, these mutations can all result in similar changes to the cells harboring them. First, they can decrease differentiation potential, compromising their use in research or therapy, or in the mouse, their ability to contribute to the developing embryo in the generation of transgenic animal models^15–18^. Second, they can confer a proliferative or survival advantage, leading to their expansion in undifferentiated cultures and posing a risk of cancer development if used for cell replacement therapy^15,18^. While around a third of human PSC samples show these structural abnormalities, many cell lines can be maintained without issue through many generations, though it is unclear why this is the case^4^. Given these problems, it is important to be able to understand what underlying variation contributes to the generation and maintenance of these aneuploidies in culture to improve our ability to propagate these valuable resources. The hypothesis that the genetic background of PSC lines might influence their genetic stability has not been investigated.

A main advantage to using mouse embryonic stem cells (mESC) as a model for studying PSC is the availability of stem cells from multiple mouse inbred strains and hybrids to facilitate genetic mapping^19,20^. However, most studies of mESC – not just in the context of aneuploidy – have had very little representative genetic diversity. One report noted variation in the frequency of specific chromosome duplications amongst cell lines from highly related genetic backgrounds^10^. Additionally, because most mESC used for research are derived from XY mice, there is little understanding of how chromosomal sex affects genetic stability in the pluripotent state. XX cells in the preimplantation epiblast, which mESC are derived from, have reactivated their second X Chromosome and therefore have higher X-linked gene dosage compared to XY mESC^21,22^. The most well documented phenotypic consequence of this dosage difference is in pluripotency signaling pathways. In serum-grown cell cultures, XX mESC exhibit a more naïve-like pluripotent state than XY mESC due to decreased MAPK/ERK signaling that presumably stems from dosage of signaling factors encoded on the X Chromosome^23^. Genetically diverse, allelically balanced multiparent mouse populations like the Diversity Outbred (DO) mouse population have been developed to perform high resolution genetic mapping projects^24^ and more recently mESC have been derived for these populations from both XX and XY mice and have been used to study genetic factors that affect maintenance of pluripotency and cell differentiation potential^20,25^. Key features of the DO panels include a high level of widely distributed genetic diversity, high heterozygosity, and accumulated recombination events that facilitate high resolution genetic mapping^26^. These features are not duplicated in existing large panels of hiPSC lines that are accrued randomly and have strong biases in their representation of genetic ancestry^27^.

While recent advances in developing chemically defined culture media have generated methods by which we can culture ESC from numerous strains, unfortunately these approaches have had adverse effects on aneuploidy frequency. Originally, mESC were cultured on a mitotically inactivated murine embryonic fibroblast (MEF) feeder cell layer which provided the mESC with the necessary paracrine signaling to proliferate and maintain pluripotency^28^. Eventually, efforts to define the culture conditions for mESC arrived at the combination of leukemia inhibitory factor (LIF) and inhibitors of the MAPK/ERK and GSK3-signaling pathways using the small molecules PD0325901 and CHIR99021 respectively, termed 2i^29^). Another, less widespread method named alternative 2i (a2i) substitutes the Src-inhibitor CGP77675 for the MAPK/ERK-signaling inhibitor^30^. While these culture conditions achieve the intended goal of a defined mESC culture media that maintains naïve pluripotency and allowed the derivation of mESC from non-permissive strains, they incidentally have detrimental effects on the genomic stability of the cells cultured in these conditions compared to conditions that include MEF feeder cells^11,31^. This highlights the need to further understand the causes of aneuploidy in mESC to develop additional media additives that can improve the maintenance of proper karyotype.

To investigate what genetic variation affects the development of aneuploidy in mESC cultures, we have used multi-omic data sets from a panel of DO mESC. These mESC were cultured in serum + LIF and only one inhibitor, the GSK3 inhibitor (for methodology, see^20^). Culturing in the presence of only this one inhibitor facilitates better understanding of the inherent differences between chromosomal sexes where most of the differences result from the MAPK/ERK-signaling pathway. Using bulk RNA-sequencing and mass spectrometry proteomics data from 170 genotyped mESC lines, we identified samples with accumulations of aneuploid cells by detecting significantly different changes in both transcript and protein expression due to changes in chromosome dosage. Analysis of these detected aneuploidies showed that cell lines with dual X chromosome dosage are less likely to be aneuploid, and a validation single-cell data set shows lower x-linked gene expression in aneuploid cells. These XX lines also have higher expression of 2C-like state genes and higher expression of tumor suppressor genes with functions in DNA damage response.

## RESULTS

### Virtual karyotyping of Diversity Outbred mESC identifies recurring duplications in four autosomes

To investigate how underlying variability in the genetics and chromosomal sex of pluripotent stem cells influences the development and accumulation of aneuploid cells, we analyzed paired-end bulk RNA-seq and multiplexed mass spectrometry proteomics data from a panel of 170 unique DO mESC lines as well as 10X single-cell RNA-seq (scRNA-seq) from an additional 12 unique DO mESC lines derived and cultured in serum + LIF/GSK3 inhibitor (CHIR99021) all collected in the Predictive Biology Inc laboratory. None of the cell lines represented in the single-cell data set are also represented in the bulk RNA-seq and proteomics data set. While the bulk data set has a nearly even number of chromosomal sexes (88 XX and 82 XY cell lines), all 12 lines in the single-cell data set are from XY embryos. After filtering out gene models, Riken genes, and genes not expressed in all mESC lines, the bulk data consists of 12,579 detected transcripts and 5,141 detected proteins. Our single-cell data consisted of 18,954 detected transcripts and 5,992 cells from between 427 to 557 per cell line.

These bulk data sets do not have associated karyotyping data, so virtual karyotyping was used to discover which chromosomes have recurrent copy number variations (CNV) accumulating within the cell line cultures. Virtual karyotyping is the process of assessing CNV in a sample by assessing the resultant increase/decrease in downstream gene expression. Our methods are similar to those implemented by other groups^6,32–35^ but specialized to suit the structure of our population and data set as well as the lack of known euploid reference control samples. Given the balanced population structure of the DO model, we expect the summary expression of each chromosome (median of gene expression for autosomes and sum of gene expression for Chromosome X) to fit a single Gaussian distribution and that a population with multiple samples harboring accumulations of CNV for that chromosome would disrupt the fit of this distribution. It is evident when looking at population-normalized gene expression across the genome that certain samples have chromosome expression levels similar to the median for the population (leftmost panels of Fig S1A) while other samples have noticeable increases in expression across entire individual chromosome(s) (rightmost panels of Fig S1A). Assaying each chromosome’s normality within the population by a Shapiro-Wilk test identified recurring duplications of Chromosomes 1, 6, 8, and 11 by their right-skewed expression distribution for both transcript and protein (Fig S1B).

Following up on the non-normally distributed chromosomes, we identified samples whose chromosome median gene expression was significantly outside of the expected distribution. After bootstrapping, we identified a total of 62/170 samples that had significant accumulations of cells with chromosome gains and none that had chromosome losses. For the purposes of brevity, the cell lines identified with this method will be referred to simply as aneuploid lines – with the implicit understanding that these are mosaics of cells with both normal and abnormal karyotype. Of the aneuploid cell lines, 10 lines (5.6%) had Chromosome 1 duplications, 8 lines (4.4%) had Chromosome 6 duplications, 35 lines (19.4%) had Chromosome 8 duplications, and 24 lines (13.3%) had Chromosome 11 duplications (Fig 1A-B). Lines with a single trisomy made up 75.81% of aneuploid samples (47/62), 78.72% of which were Chromosome 8 or 11 aneuploidies (Fig 1C). Likewise, all aneuploid lines with multiple duplications had duplication of at least Chromosome 8 or 11 in addition to another chromosome (Fig 1c).

**Figure 1.**
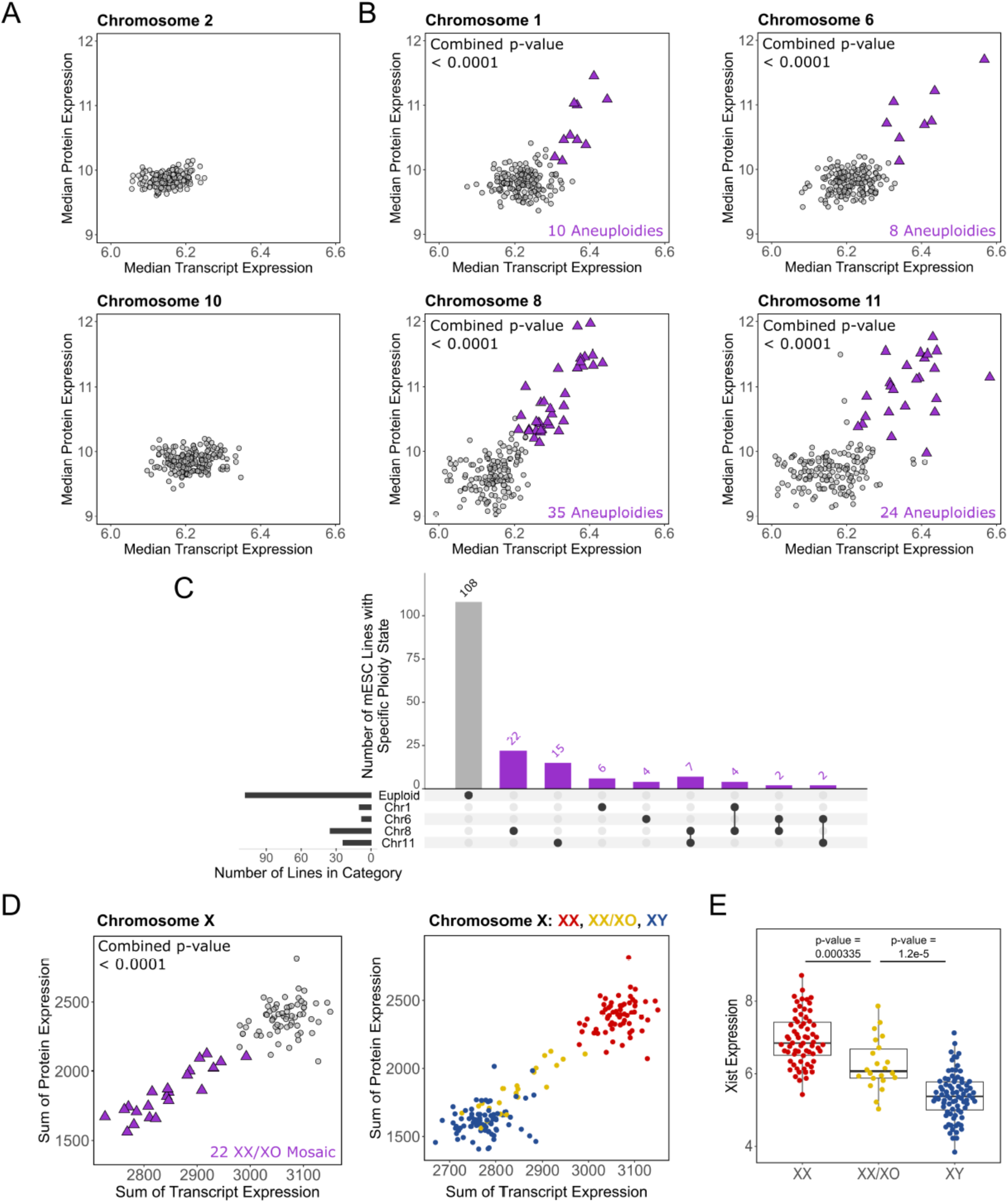
– Identification of DO mESC lines with accumulated aneuploid cell populations using RNA-seq and proteomics. (A-C) Comparison of median transcript and protein expression for all genes on specific chromosomes. (A) Commonly euploid chromosomes. (B) Commonly duplicated chromosomes. Purple triangles represent aneuploid mESC lines. (C) Number of cell lines with specific karyotypes in bulk DO data sets. (D) Comparison of median transcript and protein expression for for XX, XY, and XX/XO mosaic cell lines (E) XIST expression for XX, XY, and XX/XO mosaic cell lines from RNA-seq.

### Loss of chromosome X occurs in a subset of originally XX cell lines

Another commonly observed chromosomal copy number variation in mESC is the loss of the X Chromosome in XX cell lines^10,18^. Because both X Chromosomes are active during naïve pluripotency, we employed the same methodology that was used to determine autosomal aneuploidy to determine copy number changes in the X Chromosome for the XX lines in our data set. The expression of X-linked genes in originally XX mESC showed a left-skewed distribution, indicating that Chromosome X loss occurred in multiple DO mESC lines (Fig S1B). Bootstrapping results determined that 22/88 (25%) XX cell lines had significantly lower X chromosome expression, matching that of the XY mESC lines (Fig 1D). To rule out the possibility of improper X Chromosome inactivation (XCI), we compared the expression of Xist between high and low X expressing XX lines compared to XY male lines. The lack of substantial increase in Xist expression from the low Chromosome X gene expressing originally XX mESC lines rules out XCI, whereas the expression levels observed between these groups is more consistent with low levels of expression proportional to the number of X Chromosomes harbored by the line (Fig 1E).

### Estimating the extent of mosaicism in aneuploid cell lines

Since virtual karyotyping of bulk sequencing data alone cannot determine the percentage of aneuploid cells in a given line, we used the scRNA-seq data set to estimate the range of aneuploid mosaics we were observing. Using the R/CONICSmat package, we were able to identify individual cells with trisomies for Chromosomes 1, 6, 8, and 11 and directly compare expression of those chromosomes between euploid and aneuploid cells (Fig 2A-B). While non-duplicated chromosomes did not experience global changes in gene expression between ploidy conditions, samples with one specific trisomy had approximately a 1.4-fold increase in expression from that duplicated chromosome relative to samples with normal karyotype (Fig 2B). Knowing what chromosome-wide transcript expression change is expected if 100% of the mosaic is aneuploid for a given chromosome provides us with a way to evaluate the changes seen in the bulk RNA-seq data. Comparing each aneuploid sample to the median expression of the euploid samples, we estimate that the aneuploidies annotated in the bulk data represent between 3.44 – 9.06% cells with Chromosome 1 trisomy, 3.90 – 14.39% cells with Chromosome 6 trisomy, 2.50 – 11.63% cells with Chromosome 8 trisomy, and 2.99 – 17.27% cells with Chromosome 11 trisomy (Fig 2C).

**Figure 2.**
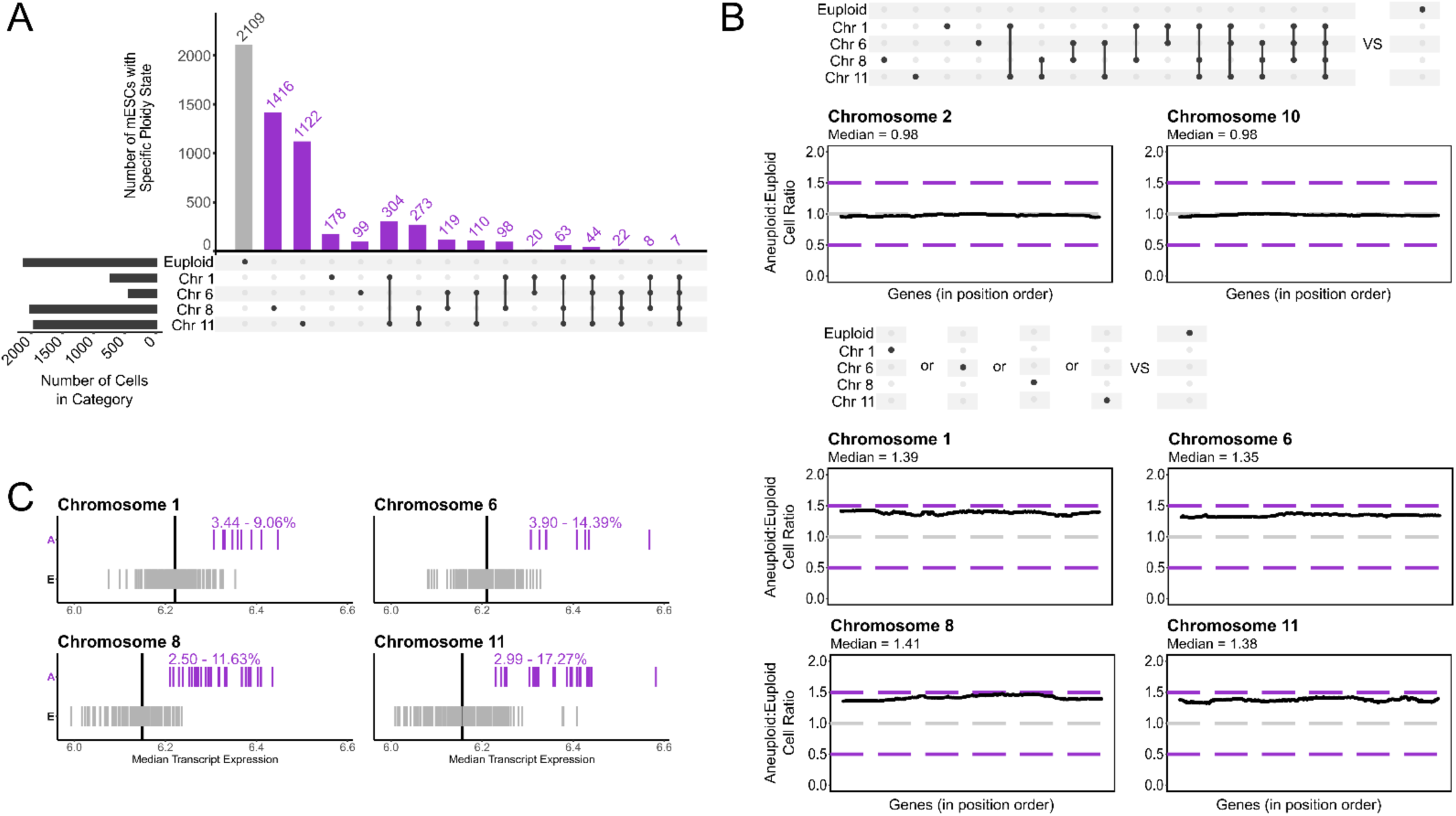
– Using DO mESC single-cell RNA-seq aneuploidy calls to estimate aneuploid content of cell lines in bulk DO data. (A) Number of cells with specific karyotypes in single-cell data. (B) Gene expression changes from euploid chromosomes in cells with abnormal karyotype compared to cells with normal karyotype (top) as well as gene expression changes from aneuploid chromosomes in cells with a specific trisomy compared to cells with normal karyotype. (C) Estimation of aneuploid cell percentage of bulk data cell lines using the changes determined from single-cell data and the median chromosome RNA-seq values from the bulk data.

### Mouse duplications replicate genes and pathways represented in human autosomal duplications

In order to understand the human relevance of the observed mouse aneuploidies, the synteny between mouse and human PSC (hPSC) duplications was assessed using The Jackson Laboratory Synteny Browser (syntenybrowser.jax.org). Genes from mouse Chromosomes 1, 6, 8, and 11 were mapped to their corresponding syntenic genes in the human genome, highlighting genes from human Chromosomes 1, 7, 12, 14, 17, 20, and X as well as the more minimal segmental duplications 1q32, 7p, 12p13.3, 17q^3^. While mouse Chromosomes 1, 6, and 11 shared broadly similar gene content to multiple human duplications, mouse Chromosome 8 was surprisingly lacking in synteny with any human duplications (Fig S2A-B). Whereas the other mESC duplications likely influence the cells in similar ways as hPSC duplications due to their shared genes, it was curious to find that the most common mouse duplication in our data had no counterpart in human aneuploid PSC. For this reason, we wondered if the trisomy 8 mESC might alter the same pathways as human chromosomal duplications through a different subset of genes.

To identify the consequences of trisomy 8, differential expression analysis was performed on the scRNA-seq data set because it represents purer euploid and aneuploid populations than the bulk data. All cells harboring only trisomy 8 were compared to cells with no detected aneuploidies. As expected, the comparison showed differential expression of pathways related to gene expression/processing as expected when accommodating a large-scale increase in genetic material (Fig S2C). The trisomy 8 cells also differentially expressed genes related to the DNA replication, consistent with the observation that aneuploidies can drive a proliferative advantage relative to euploid cells (Fig S2C). Most of the downregulated pathway terms related to immune response (Fig S2C). Despite a paucity of shared genes with human PSC mouse trisomy 8 seems to similarly result in alterations to gene expression and processing mechanisms and cell proliferation, similar to hPSC.

### Loci on Chromosome X associated with overall autosomal ploidy

We began interrogating variation that affected autosomal aneuploidy in the DO samples by mapping genotypes associated with aneuploidy status. Genetic mapping was performed using the R/qtl2 package, using aneuploid and euploid status as a binary trait while correcting the covariates to reflect the recently identified XO lines previously labeled as XX. No significant loci were discovered for any individual trisomy (Fig 3A), however a significant locus at Chromosome X was discovered for general abnormal karyotype (p < 0.01; Fig 3B). The founder effects under this interval indicate WSB/PhJ alleles associated with higher likelihood of aneuploidy and PWK/PhJ alleles associated with lower likelihood of aneuploidy (Fig 3C). Unfortunately, genes under this interval did not immediately stand out as good candidates. Therefore, we believe it is more likely that genetic variation in this region affects aneuploidy by affecting broader changes, such as structural variants might cause.

**Figure 3.**
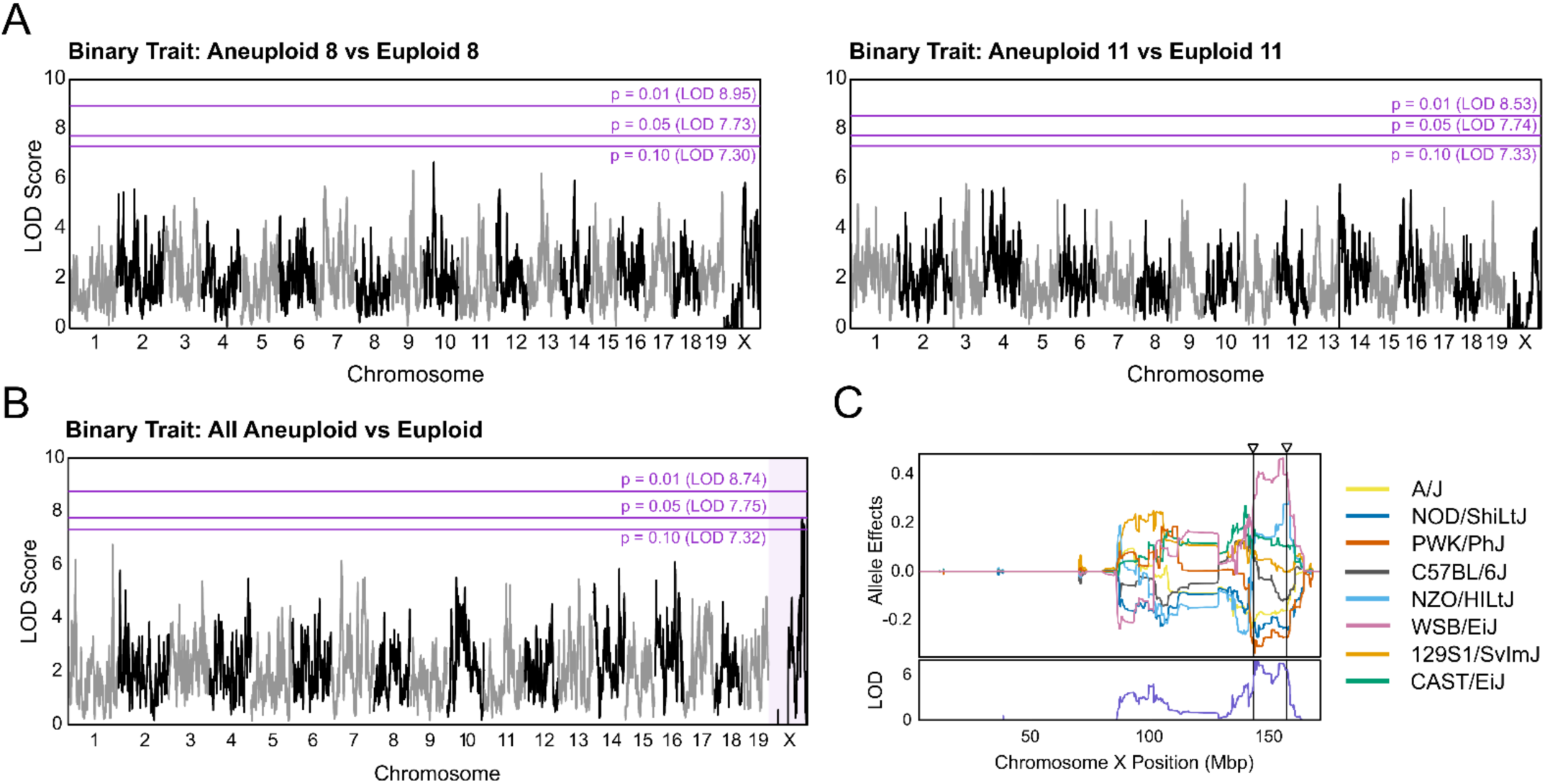
– Genomic mapping of DO mESCs using aneuploidy as a binary trait reveals influence of Chromosome X variation. Assessing the relationship between genomic loci with duplication of Chromosome 8 or 11 (A) and any autosomal duplication (B). (C) Effects of alleles from DO founder mouse strains on aneuploidy at the significant locus found on Chromosome X.

### Cell lines with two active X chromosomes have lower aneuploid proportions

Following from the observation that genetic variation on Chromosome X was associated with abnormal karyotype, we assessed the proportion of aneuploid mESC lines within each chromosomal sex. There was a significantly lower proportion of aneuploid XX lines compared to XY or XO lines (11/66 XX lines, 41/82 XY lines, 10/22 XO lines; Chi-squared p-value = 0.0001; Fig 4A). This pattern was not only observed with overall karyotype, but also for samples with trisomy 8 and 11 (4 and 3/66 XX lines, 26 and 16/82 XY lines, 5 and 5/21 XO lines; Chi-squared p-value = 0.0006 & 0.0158 respectively; Fig 4A). Although too few samples were annotated as trisomy 1 or 6 to detect significance, the same trend between XX and XY or XO samples can be seen for these abnormalities as well (Fig 4A). Except for trisomy 11, the proportion of aneuploid XO lines fell in between the XX and XY samples, which suggested that X Chromosome dosage was underlying the observed stratification since the mosaic of XO samples includes some proportion of XX cells as well.

**Figure 4.**
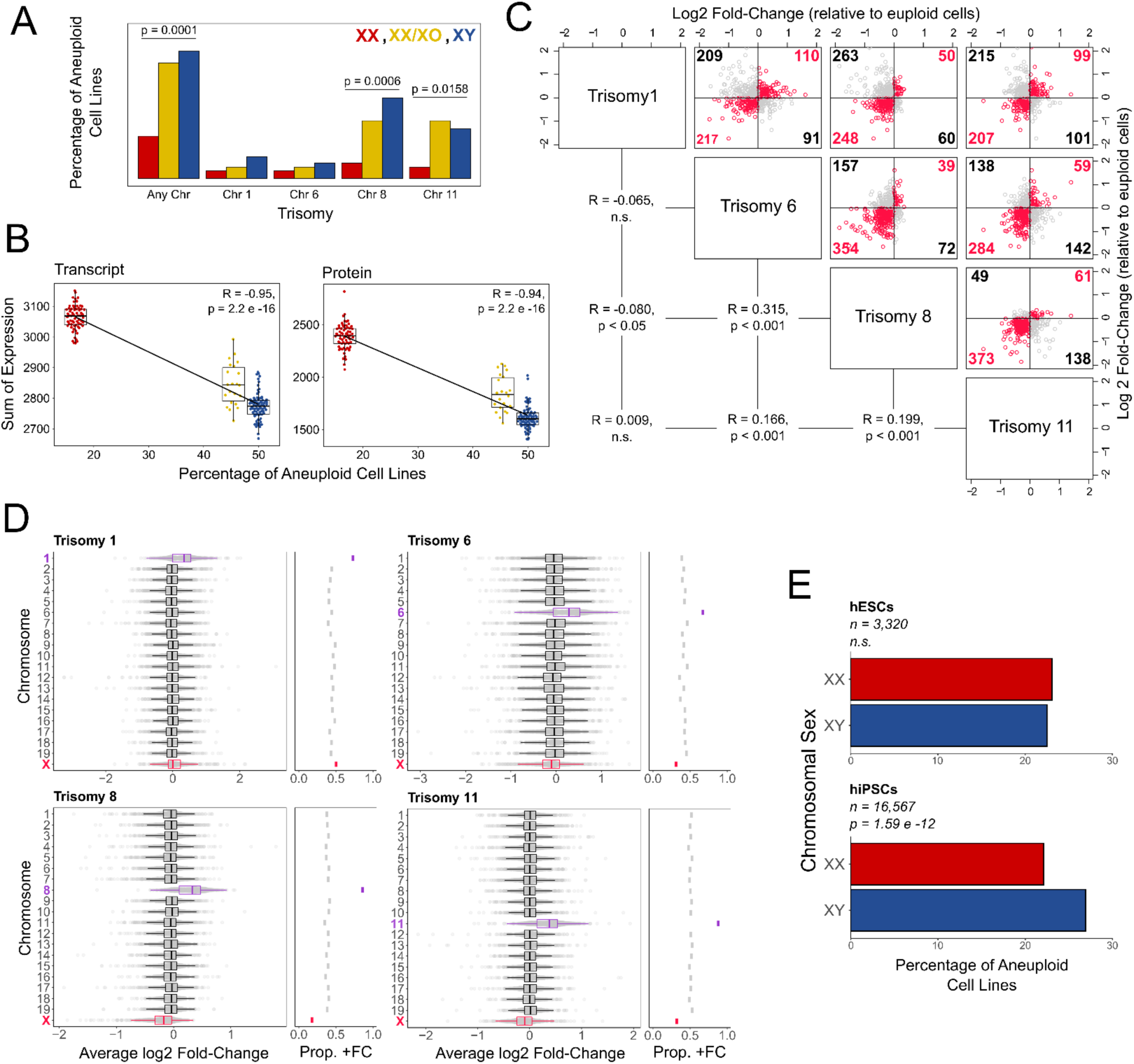
– Inverse relationship between Chromosome X gene dosage and autosomal duplications. (A) Proportion of cell lines from the bulk DO data sets classified as aneuploid by Chromosome X dosage: two (XX), mosaic of one and two (XX/XO), and one (XY). (B) Correlation between each chromosomal sex’s expression of Chromosome X transcripts or proteins and the proportion of aneuploid cell lines using DO bulk data. (C) From scRNA-seq of DO mESC lines, the correlations between X-linked gene expression changes in cells with a specific trisomy (relative to euploid cells). Red points are genes with shared directional changes – increasing in both trisomies or decreasing in both trisomies. (D) Average gene expression changes in cells with a specific trisomy and the proportion of genes with positive changes in each chromosome from the single-cell DO data. The duplicated chromosome is highlighted in purple and Chromosome X is highlighted in red. (E) Percentage of XX and XY hiPSC and hESC cell lines submitted to WiCell with identified autosomal duplications by G-banded karyotyping.

To see if there was a relationship between expression of the X Chromosome and likelihood of being annotated as aneuploid, we assessed the correlation between global Chromosome X gene expression and proportion of aneuploid lines within each chromosomal sex. For global Chromosome X expression, the same sum of transcript or protein expression from Fig S1 & Fig 2 were used. There was a distinct negative correlation between expression of X-linked genes and the proportion of aneuploid lines in a sample (Pearson correlation = −0.95, p = 2.2×10^−16^ for transcripts and Pearson correlation = −0.94, p = 2.2×10^−16^ for protein; Fig 4B). This observation served as additional evidence that the aneuploid-associated loci on Chromosome X reflect X-linked gene expression more broadly rather than a single causal gene under the interval.

Using the aneuploidy annotations in the scRNA-seq data set of all XY mESC lines, we were able to observe a similar pattern at the individual cell level. We performed differential expression analysis comparing cells with a single trisomy to completely euploid cells and analyzed the proportion of genes from each chromosome that were upregulated, downregulated, or experienced no changes. As expected, there was a large proportion of upregulated genes from the duplicated chromosome for each individual trisomy (Fig 4C). The remainder of the autosomes shared a similar ratio of up- and down-regulated genes across trisomies, however in all besides trisomy 1 there is a higher proportion of downregulated Chromosome X genes (Fig 4C). The log2 fold-change in expression of X-linked genes was also significantly correlated between trisomy 6, 8, and 11 (0.315 between 6 and 8, 0.166 between 6 and 11, 0.199 between 8 and 11; all p < 0.001; Fig 4D). These results demonstrate that individual aneuploid cells, even within all XY cell lines, have lower X-linked gene expression. Because the scRNA-seq data is at a single time-point, we cannot determine whether individual cells with lower Chromosome X expression are more likely to develop aneuploidies (as observed in the bulk data) or if this is evidence that aneuploidy induces decreased expression of genes on Chromosome X.

We were interested to see if human PSC shared the observation from mouse models that Chromosome X dosage modified the likelihood of aneuploidy. Collaborators at The WiCell Institute and Sheffield University analyzed the human ESC and iPSC lines submitted to The WiCell Institute for karyotyping. Although the differences were less pronounced than in the mESC, there was a general trend of increased abnormal karyotype in the XY cell lines compared to the XX lines in iPSCs, but not in ESCs (p = 1.59×10^−12^ in iPSCs; Fig 4E). Although the effect sizes are different, we do not expect identical effects due to species differences in culture conditions and chromosome X dynamics during cell culture. Interestingly, observing this aneuploidy difference in iPSCs and not in ESCs runs contrary to expectations based on the state of Chromosome X erosion in these two cell types reported in^36^, although it be the result of which genes are reactivated and at what time they are reactivated during erosion. This data from thousands of directly karyotyped cell lines by an external source are strong evidence that this is not a mouse-specific relationship, although the additional nuances of human PSC culture may mitigate this effect under some conditions.

### XX mESC cultures accumulate less aneuploidy over time in culture than isogenic XY mESCs

To validate our retrospective observation that fewer aneuploid XX mESC lines were identified than XY mESC lines in a different cellular system, we performed scRNA-seq on isogenic C3H/HeSn-Paf/J XX, XY, and XO mESC lines at the beginning and end of ten passages in Serum/LIF culture. For each chromosomal sex, three independently derived cell lines were aged for the study. After cells were demultiplexed and filtered, our early isogenic scRNA-seq data set consisted of 5,200 XX cells, 5,375 XY cells, and 3,564 XO cells and our late scRNA-seq data set consisted of 2,832 XX cells, 5,511 XY cells, and 16,600 XO cells (Fig S3, Fig S4). These early and aged data sets represented expression of 25,573 and 25,452 genes respectively. The large increase in XO cells indicates that some of the XX and XY cells lost their second sex chromosome over time, although we cannot tell for certain because the demultiplexing was performed based on transcriptomic signatures of each chromosomal sex and not by external methods.

First, we annotated the autosomal aneuploidies present at both timepoints to compare with the DO mESC data sets (Fig 5A). Our early timepoint began at 86.6% entirely euploid cells, which decreased to 19.7% by the end of ten passages. The population of aneuploidies in the early timepoint was predominantly Chromosome 8 and 11 trisomies, matching the distribution observed in the DO mESC bulk and single-cell data. However, the aged aneuploid cells were mostly Chromosome 1 and 8 trisomies – some of which harbored additional chromosome duplications. This observation suggests that while duplication of Chromosome 11 is more common in earlier cultures, Chromosome 1 duplication may provide advantages that prevail over long periods of time. Another interesting observation was the identification of two duplications not seen in the DO data sets: duplication of Chromosomes 14 and 19. These duplications were observed almost exclusively in conjunction with other previously noted aneuploidies (mostly Chromosome 1) and have not been identified as commonly as others in the literature. Therefore, we suspect that they are secondary aneuploidies that provide an advantage in culture when in the presence of other acquired advantages from more common duplications.

**Figure 5.**
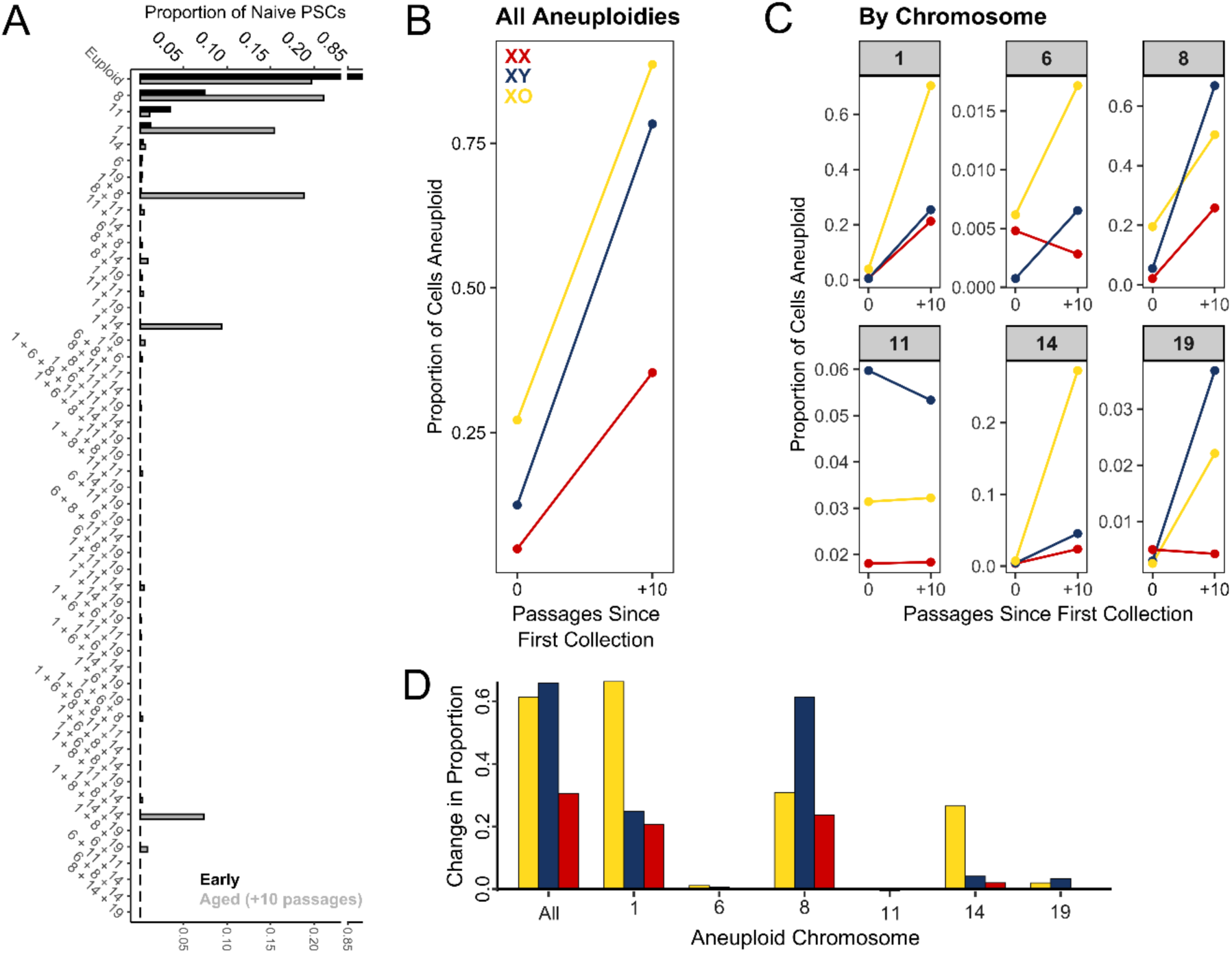
– In vitro aging of isogenic cell lines with different chromosomal sexes. (A) Proportions of specific chromosomal duplications in early and late (aged) passage mESCs identified from single-cell RNA-seq. (B-C) Before and after proportions of (B) total aneuploidy and (C) specific chromosomal duplications in each chromosomal sex. (D) Change in proportion of total aneuploidy and specific chromosomal duplications in each chromosomal sex.

Importantly, analysis of the aneuploid annotations in these aged isogenic mESC also demonstrated a larger increase in aneuploid cell population in the XY and XO lines compared to the XX lines. While XX lines already had fewer aneuploid cells at the early collection, they also had a smaller change in aneuploid proportion over the course of ten passages than the XY and XO lines had (Fig 5B,C,D). When broken down by specific trisomy observed, the differences due to chromosomal sex were less distinct (Fig 5C). However, this is not unexpected if we assume that selection for specific chromosomal duplications occurs only after missegregation. Therefore, different cultures may accumulate different aneuploidies based on which chromosomes are randomly allocated to the wrong daughter cell. An alternate hypothesis could be that specific aneuploidies are differentially beneficial to cells with different suites of sex chromosomes. Overall, the data shows that the X Chromosome dosage of a cell line influences the buildup of aneuploid populations in culture, validating the observation from diverse mESC.

### Dusp9 expression correlates with mESC sexual dimorphisms and ploidy status

While it may not be likely that a single causal gene is responsible for the differential aneuploidy proportions between single and dual X Chromosome samples, we looked for highly associated genes to better understand the molecular underpinnings of this effect. Because XX lines are seldom used in mESC studies, the literature on their sexual dimorphisms is sparse. However, one review noted a difference in the pluripotency signaling pathways between XX and XY mESC under certain culture conditions^37^. The review identified six X-linked genes (*Eras*, *Nono*, *Dusp9*, *Tfe3*, *Nr0b1*/*Dax1*, and *Zfx*) which were associated with pluripotency and able to at least partially replicate the differences in molecular markers of pluripotency (increased *Nanog*, *Esrrb*, and *Tcl1* expression, Erk phosphorylation, and Akt activity; decreased Mek, Erk, and Gsk3 expression, and DNA methylation; prevention of differentiation) – of which *Dusp9* overexpression was most similar to increased X dosage.

Because protein expression is more highly attenuated than transcript expression, we calculated the transcript∼protein correlation of X-linked genes in our bulk RNA-seq and proteomics data to better understand which genes are most affected by increases in X dosage at the functional level – paying attention to the genes identified in^37^. *Dusp9* had the highest transcript∼protein correlation (0.8207) of all six^37^ genes, the next highest of which was *Eras* (0.6361; Fig 6A-B). Within each sex chromosome category, there is lower expression of *Dusp9* transcript and protein in aneuploid lines compared to euploid lines (Fig 6B-C). Therefore, *Dusp9* best phenocopies the pluripotency differences conferred by increased X dosage^37^ is highly dosage sensitive – being more lowly expressed in more aneuploidy-prone single X lines, and is also more lowly expressed in those lines that have accumulated aneuploid cells.

**Figure 6.**
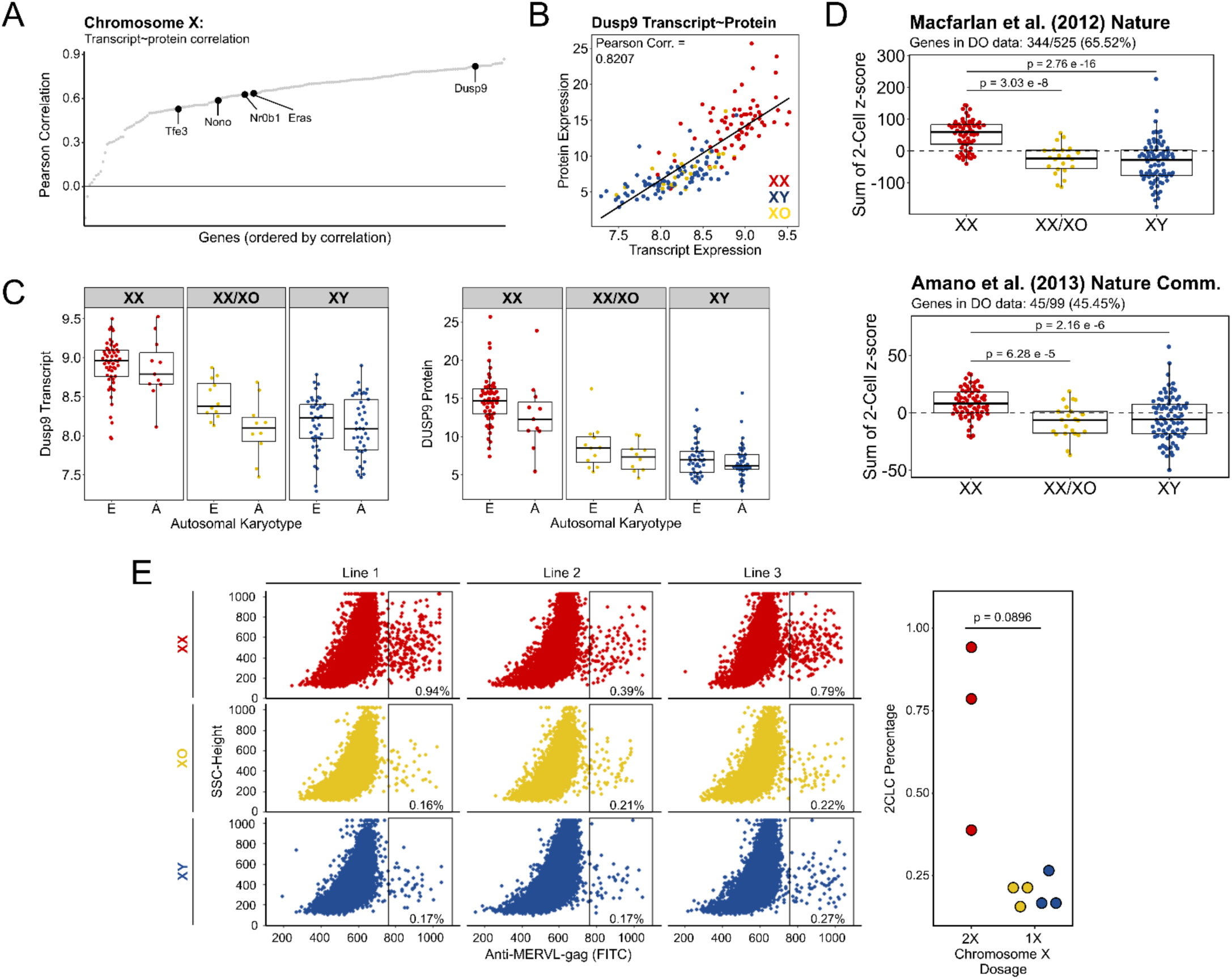
– Increased proportion of 2C-like state cells in mESCs with two X Chromosomes. (A) Dose sensitivity measured by the correlation between transcript and protein for all X-linked genes represented in the RNA-seq and proteomics in the DO bulk data. Genes that may account for the pluripotency state differences between XX and XY mESCs are labeled^23^. (B) Comparison of Dusp9 transcript and protein abundance in individual DO mESC lines colored by chromosomal sex. (C) Abundance of Dusp9 transcript and protein in individual DO mESC lines, comparing euploid and aneuploid lines within each chromosomal sex category. (D) Abundance of transcripts upregulated in the 2C-like state as identified in^39,40^ within each chromosomal sex in bulk RNA-seq data. (E) Flow cytometry for the 2C-like state marker MERVL gag protein in XX, XO, and XY mESC lines from the same genetic background. Signal from each cell (left) and percent of cells with increased MERVL-gag expression from each line (right).

### Cell lines with two active X chromosomes have higher expression of genes upregulated in 2CL state

Differential *Dusp9* expression led us to the consideration of a rare, metastable state recently discovered in mESC called the 2-cell-like (2CL) state. This is a state that mESC periodically transition in and out of during culture, and while generally less than one percent of a culture will be in the 2CL state at any given time depending on conditions and cell line, within about nine passages all of the cells will have gone through that transition^38^. The 2CL state is named for the resemblance of its transcriptional profile to the 2-cell stage of embryonic development, particularly the expression of *Zscan4* and endogenous retroviruses like murine endogenous retrovirus L (MERVL)^39,40^. Zscan4 functions in Tert-independent telomere elongation^38^ and according to^41^ may either be involved in supernumerary chromosome removal or may bias survival in favor of normal karyotypes. The *Dusp9* expression patterns that we observed were of interest because feeder-based mESC culture conditions not only favor genomic stability, but also promote an increased presence of 2CL state cells and show strong upregulation of *Dusp9* compared to feeder-free culture^11^. Since *Dusp9* is expressed higher in dual X mESC lines, euploid mESC lines, as well as feeder-based mESC cultures and all of those are associated with improved genomic integrity, it may be that a common thread is the 2CL state.

In order to better understand the relationship between X Chromosome dosage and the 2CL state, we compared expression of 2CL state genes between sex chromosome categories. Lists of transcripts upregulated in 2CL state cells were obtained from^39,40^. Within our DO mESC bulk RNA-seq data, we found matches for 344/525 (65.52%) of the Macfarlan genes and 45/99 (45.45%) of the Amano genes. The sum of the 2CL state gene expression z-scores were significantly higher in XX mESC lines compared to the XY or XO mESC lines in both gene sets used, suggesting either a stronger induction of these genes or a higher proportion of 2CL cells (2CLCs) in the XX mESC (Fig 6D). To investigate the 2CL differences, isogenic mESC from XX, XO, and XY lines were assayed by flow cytometry against MERVL gag protein to quantify the number of 2CLCs in each chromosomal sex. While the 2CLC population was small in each cell line as expected, the number of 2CLCs was consistently higher in XX mESC compared to the XO or XY mESC, consistent with the observations from bulk RNA-seq data (Fig 6E). These results combined with the chromosomal sex bias in aneuploidy development suggest that an increased dosage of X-linked genes in XX mESC increases the proportion of 2CLCs which may account for the improved genomic stability of these cell lines.

### Aneuploid mESCs Disregulate X-Linked Tumor Suppressor Genes

Because the human PSC subpopulation comparable to the 2C state, the 8-cell-like (8CL) state, has not been reported in the primed state culture conditions that most of the WiCell data comes from, we sought an alternative explanation for the shared effect of Chromosome X dosage between species. We hypothesized that X-linked tumor suppressor genes (TSG) may be differentially expressed in XX cell lines, therefore counterbalancing the selective advantages that autosomal aneuploidies may confer. We collected a list of 24 TSG on Chromosome X from the Tumor Suppressor Gene Database (bioinfo.uth.edu/TSGene) and analyzed their dosage-sensitivity in our bulk data sets. *Dusp9* also came up in our list of TSG, but two other genes had notable dosage sensitivity: *FHL1* and *KDM6A* (Fig 7A). Of these, only *FHL1* was significantly associated with ploidy status as well as Chromosome X dosage (Fig 7B).

**Figure 7.**
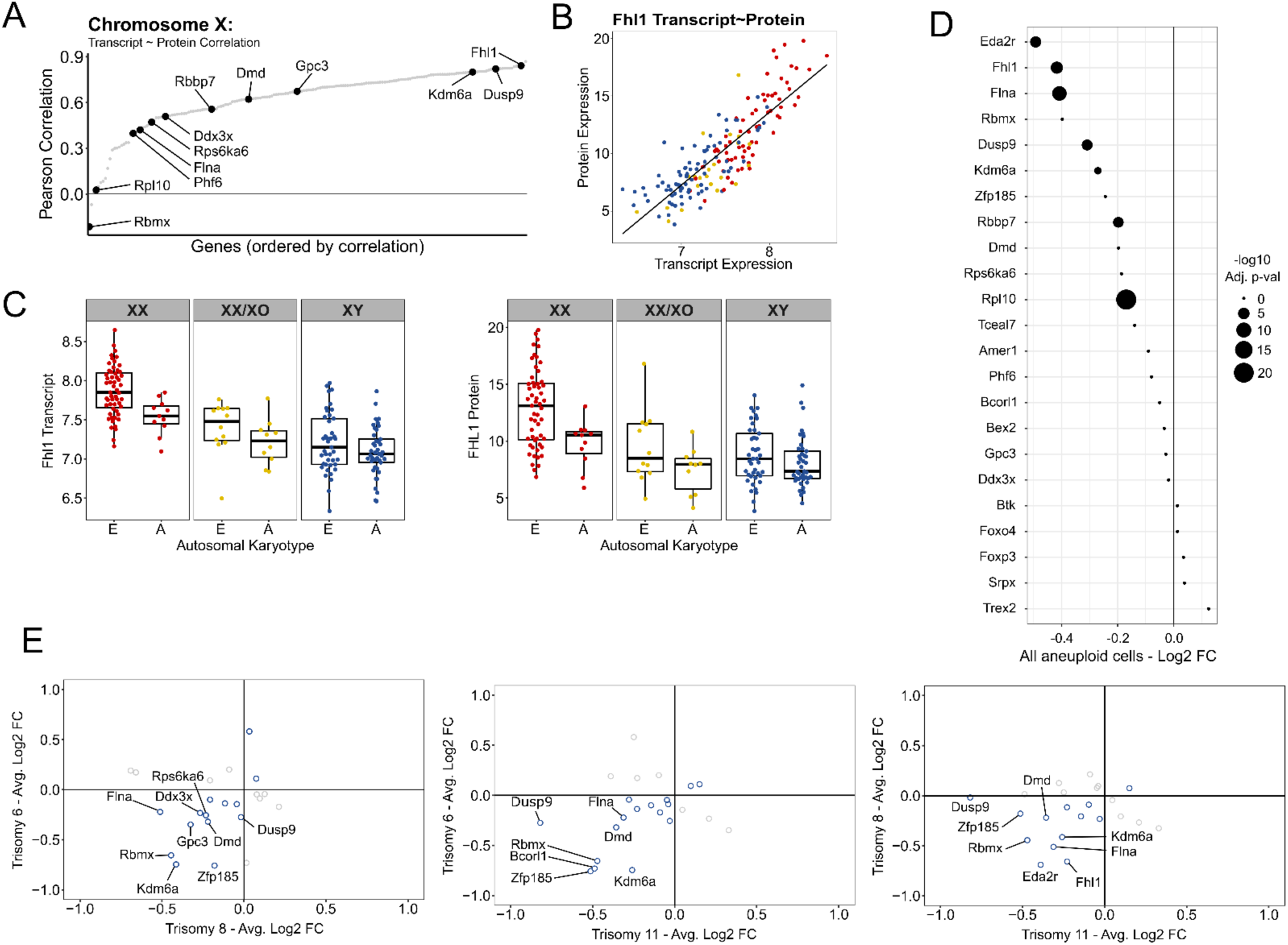
– Varied modulation of X-linked tumor suppressor genes in aneuploid mESCs. (A) Dose sensitivity measured by the correlation between transcript and protein for all X-linked genes represented in the RNA-seq and proteomics in the DO bulk data. Genes identified as tumor suppressor genes by the Tumor Suppressor Gene Database are labeled (bioinfo.uth.edu/TSGene). (B) Comparison of Fhl1 transcript and the most predominant protein isoform abundance in individual DO mESC lines colored by chromosomal sex. (C) Abundance of Fhl1 transcript and the most predominant protein isoform in individual DO mESC lines, comparing euploid and aneuploid lines within each chromosomal sex category. (D) Changes in transcript expression of X-linked tumor suppressor genes between cells with abnormal compared to cells with normal karyotype in DO scRNA-seq. (E) Changes in transcript expression of X-linked tumor suppressor genes between cells with a specific trisomy to cells with normal karyotype in DO scRNA-seq. Tumor suppressor genes with the most notable changes are labeled.

Additionally, because we noted that aneuploid XY mESCs are associated with lower Chromosome X gene expression, we wanted to assess changes to these X-linked TSG specifically. When observing log2-fold changes in all aneuploid cells, there were clear decreases in expression of *Eda2r*, *Fhl1*, *Flna*, *Rbmx*, *Dusp9*, *Kdm6a*, *Zfp185*, *Rbbp7*, *Dmd*, *Rps6ka6*, and *Rpl10* – of which the three dosage sensitive genes we identified were in the top six (Fig 7C). However, when comparing specific aneuploidies due to their unequal representation in the scRNA-seq data set, it becomes apparent that these TSG have variable differential expression between trisomies (Fig 7D). These data suggest that some TSG are expressed higher in cells with two Chromosome X copies and no one X-linked TSG may be responsible for the relationship between their expression and aneuploidy.

## DISCUSSION

Here we have presented a methodology to virtually karyotype a diverse group of cell lines without euploid controls by using both transcriptomic and proteomic data. We were able to show patterns of aneuploidy that were similar to that found by an established R package for karyotyping single-cell data sets (Fig 1, Fig 2, Fig 5). These patterns showed that the DO mESCs did not sustain any losses of autosomes, only losing Chromosome X in a subset of the XX cell lines (Fig 1C-D). Of the duplications observed, Chromosomes 1, 6, 8, and 11 were recurrent in both our bulk and single-cell data sets, although there was a distinctly higher likelihood of Chromosomes 8 and 11 being duplicated (Fig 1B, E, Fig 2A). While we have tried to make these results generalizable by using a panel of cell lines with a highly diverse genetic background and even split of chromosomal sexes, there is still the possibility that trends in CNV may also reflect the culture conditions and variations therein may favor increased/decreased dosage of different gene sets. However, it seems that on the whole, the duplications we commonly see in mouse and human generally share duplications of the same genes or at least similar pathways (Fig S2), with the notable exception of chromosome X which is lost in mouse PSC cultures and gained in human PSC cultures^7^.

Our results reveal a distinct connection between the expression of Chromosome X genes and the maintenance of normal karyotype in mESC. There are clear examples of the direct association of lower Chromosome X dosage with increased likelihood for aneuploidy (Fig 4A-B, Fig 5). Moreover, aneuploid cells with the same number of X Chromosomes as their euploid counterparts in the single-cell data set show statistically lower expression of Chromosome X genes (Fig 4C-D). There is evidence in models outside the PSC field to suggest that there may be a bidirectional relationship between X-linked genes and genomic stability in multiple species. Individuals with Turner Syndrome (XO) have an increased risk for solid tumors compared to XX individuals, while those with Klinefelter Syndrome (XXY) have reduced solid tumor risk compared to XY individuals^42^. Likewise, there is a noted male bias in the development of many types of cancer^43–46^. In a follow up on this observation, another lab discovered a reduced ratio of X to autosomal gene expression in a number of tumor samples^47^. There have also been experiments in Drosophila demonstrating that induced autosomal CNVs result in reduced expression of Chromosome X, similar to what we observed in the single-cell data ^48^. The mechanisms that underpin each directionality of this relationship might still be distinct. Certain genes on Chromosome X may promote improved genomic stability, however the expression of X-linked genes – including those promoting genomic stability – may be decreased as a consequence of autosomal duplications. These observations mostly come from cancer models, and to the best of our knowledge, this is the first observation of chromosomal sex bias in the genomic integrity of ESC.

Our observations on X Chromosome dosage and aneuploidy in mESC were reflected in a large cohort of hiPSC as well, though the effect size was smaller (Fig 4E). A differing impact of X dosage in the two species is not a wholly surprising result given the different and complex dynamics of the X Chromosome during early development, and the variations across in vitro culture models between and within each species. While mESC seem to consistently reactivate their second Chromosome X during culture, there are variable states of reactivation during the culture of human ESC and iPSC. The most widely used human culture models appear to be more akin to the primed pluripotent state where XCI has taken place^49,50^. Additionally, human ESC and iPSCs have different predisposition for the erosion of gene expression from the X Chromosome, dampening expression from genes in the interior of Chromosome X even during reactivation^36^. Overall, XX hPSCs are less likely to express two full X Chromosomes than their mouse counterparts due to XCI or incomplete XCI erosion. Therefore, it would be expected that this effect would be diminished in an hPSC model, however it suggests that genes which escape XCI may be crucial players in the prevention of chromosomal CNV as has been suggested in the cancer field^51^.

The story we presented led to the novel discovery that cell lines with two active X Chromosomes have an increased proportion of cells in the rare 2CL state at a given time (Fig 6D-E). There have been studies showing that fostering the 2CL state benefits the genomic stability of mESC, possibly through enhanced telomere maintenance^11^. Another study hinted at a restorative role for the 2C state, noting the emergence of a population of euploid cells in a culture of cells derived from trisomic donors through overexpression of the 2C gene Zscan4^39^. The observation that XX mESC lines have higher 2CL proportion and less likelihood to develop aneuploid cells than XY or XO lines is a strong indication that the two are related, however this is currently correlative and not causative. Future study connecting differential 2CL state and genomic stability in this model must be conducted to show clearly that increased 2CLC populations in XX mESC lines are sufficient to decrease aneuploidy development. Efforts to induce a long-term increase in the 2C proportion of cells in our cultures resulted in instability after weeks of culture (not shown).

During the conception and majority of the execution of this project, the presence of a totipotent state in PSC culture was considered a mouse-specific phenomenon. As of writing, several papers have been published on the existence of 8-cell-like cells (8CLC) within human naïve PSC cultures^52–55^. These 8CLC have similar transcriptomes to the 2CLC of mouse cultures, expressing totipotency genes such as Zscan family genes and endogenous retroviruses^52,53^. These subpopulations appear in small quantities – similar to the 2CLC in mouse – when using culture media designed to promote naïve PSC characteristics, such as PXGL, 5iLA, 5iLAF, t2iLGö, RSeT, but interestingly they were not seen in expanded potential stem cells^52,53^. It is not known whether naïve PSCs bearing two X Chromosomes would be more stable genetically, or if they would show a higher proportion of 8C-like cells (Fig 6E).

Because the dosage of Chromosome X effects both mouse and human PSC, albeit to different extents based on the data here, the explanation for these effects is likely in part a shared mechanism. Since no evidence for the 8CL state in human primed PSC exists, we proposed that this shared mechanism was the increased dosage of TSG expressed from Chromosome X, as there is evidence that some escape XCI^51^. We do not notice a shared pattern of which TSG are correlated with euploid or aneuploid cells. While *Dusp9* and *Fhl1* stand out from the bulk data, the scRNA-seq data show a more trisomy-specific suite of TSG with higher expression in euploid single cells (Fig 7). These results are not indicative of a lack of TSG involvement, but rather that no one specific TSG is responsible for the aneuploidy mitigation from Chromosome X dosage.

While the increase in the proportion of 2CL state cells may be a reason for the aneuploidy differences between mESC with different numbers of Chromosome X copies, it may only be part of the disparity. As previously mentioned, mouse PSC and human PSC are not typically maintained in exactly the same naïve state during in vitro culture and evidence of a totipotent population has not been observed in the most predominant human culture systems that represent the majority of data from WiCell. Human XX naïve PSC show variability in the activation of the X Chromosome with varying patterns of erosion^56^. Existing datasets on naïve hPSC are not large enough to enable genetic analysis of chromosomal stability (Nissim Benvenisty, personal communication). Thus, the relationship between X dosage, the 8C-like state, and genetic stability in naïve PSC remains uncertain.

Our results have implications for prevention of recurrent genetic abnormalities in hPSC. The finding of synteny between affected genomic regions in mESC and hPSC suggests common underlying selective mechanisms and validates the mESC as a model for human cells. The lack of evidence for specific autosomal loci with a strong influence on development of aneuploidy suggests that variations in cell culture environment may be critical to the variable occurrence of aneuploidy across cell lines and cell labs. This is in accord with recent observations from Barbaric and co-workers^57^. Lastly it has long been known in human that males are more susceptible to cancer development than females. Our work suggests that the mESC system might be a powerful tool to investigate the underlying mechanisms of this sex-dependent effect, particularly as aneuploidy is strongly associated with progression of malignancy in mouse pluripotent stem cells^58^.

## EXPERIMENTAL PROCEDURES

### Diversity Outbred mESCs

The DO mESC lines used in this study were the same lines published in^20,59^ and derived, cultured, and harvested as described in those methods sections. Briefly, DO mice were purchased from The Jackson Laboratory and bred at Predictive Biology Inc. The ICM from the resulting embryos were prepared as described in^19^ to derive mESC. Following derivation in serum + 2i/LIF media, mESC were transferred to a serum + 1i/LIF media (GSK3 inhibitor CHIR99021 only) and slowly tapered off feeder-based culture until collection for bulk or scRNA-seq and mass spectrometry proteomics.

### Diversity Outbred mESC Bulk RNA-seq

RNA-seq data was collected, assayed, and processed as described in^20,59^ with the following exceptions. Whereas ^20^uses the single-end version of this data set, we use the paired-end data used in^59^. Both previous papers used reads that had been rank Z transformed, but here we omitted this step as this transformation removes the inherent structure necessary to identify copy number variations from non-normally distributed expression values. Before analyses, genes not expressed in all cell lines, gene models, and Riken genes were excluded.

### Diversity Outbred mESC Proteomics

Mass spectrometry proteomics data was collected and assayed as described in^59^ and processed as in^60^ with the following exceptions. Like the bulk RNA-seq data, these data have not been rank Z transformed in order to preserve the structure necessary to identify copy number variations from non-normally distributed expression values. Before analyses, genes not expressed in all cell lines, gene models, and Riken genes were excluded.

### Diversity Outbred mESC Single-cell RNA-seq

DO mESC 10X Genomics scRNA-seq gene x cell matrix was provided by Ted Choi of predictive Biology. Data processing was performed mostly as described in the standard R/Seurat package tutorial. Only cells with less than 10% of total reads from mitochondrial genes, greater than 2,000 detected genes, and that had unique cell line identification were kept for further analysis. Cell cycle scoring was performed using mouse cell cycle genes from https://raw.githubusercontent.com/hbc/tinyatlas/master/cell_cycle/Mus_musculus.csv, but no significant effects were identified that would necessitate regression.

### Isogenic mESC Single-cell RNA-seq

For sequencing, cells were dissociated as they were for flow cytometry (see below for described methods). Once lifted, cells were allowed to settle in a 10cm culture dish to remove MEF feeders due to the differential attachment rate of mESC and feeders. The remaining cells were then passed through a 40 micron filter to remove clumping cells. Then, 500,000 cells from each cell line were labeled with lipid modified oligos (LMO) as described in^61^. Briefly, cells were washed and resuspended in ice-cold PBS before incubating with 1:1 molar ratio of barcode oligo and LMO anchor (supplied by Chris Baker from the lab of Chris McGinnis) for 5 minutes at room temperature. Then, cells were incubated with LMO co-anchor (also from Chris Baker) on ice for 5 minutes. LMOs were blocked with 1% BSA and 1% FBS in ice-cold PBS, washed in the same solution, and sent for 10X library prep and sequencing in The Jackson Laboratory single-cell biology core facility.

Cell viability was assessed on a Luna FX7 automated cell counter (Logos Biosystems), and up to 40,000 cells (4,000 cells per LMO) were loaded onto one lane of a 10X Chromium Controller. Single cell capture, barcoding and library preparation were performed using the 10X Chromium platform^62^ version 3.1 NEXTGEM chemistry and according to the manufacturer’s protocol (#CG00315) with modifications for generating the LMO libraries. cDNA and libraries were checked for quality on Agilent 4200 Tapestation, quantified by KAPA qPCR, and libraries were pooled using a ratio of 95% gene expression library and 5% LMO library before sequencing; each gene expression-LMO library pair was sequenced at 91% of an Illumina NovaSeq 6000 S4 flow cell lane, targeting 20,000 barcoded cells with an average sequencing depth of 100,000 read pairs per cell.

Illumina base call files for all libraries were converted to FASTQ using bcl2fastq v2.20.0.422 (Illumina) and FASTQ files associated with the gene expression libraries were aligned to the GRCm38.93 reference assembly with GENCODE vM23 annotations (10x Genomics mm10 reference 2020-A) using the version 6.1.2 Cell Ranger count pipeline (10x Genomics). FASTQ files representing the LMO libraries were processed into cell x LMO digital count matrices using CITE-Seq-Count (version 1.4.5; https://zenodo.org/badge/latestdoi/99617772).

### Computational Analyses Virtual Karyotyping of Bulk Transcriptomics and Proteomics

Virtual karyotyping of diversity outbred mESC bulk data was performed on batch corrected expression data that had not been rank Z transformed. Gene models and Riken genes were removed from the data set before analysis. Chromosome summary statistics for each sample were generated by calculating the median of expression values for all genes on autosomes and sum of expression values for Chromosome X. Chromosomes with copy number variations were identified by Shapiro-Wilk normality test, where chromosomes determined non-normal by p ≤ 0.01 for RNA and protein were determined to represent reoccurring CNV. Specific samples harboring these abnormalities were identified by bootstrapping gene expression values to calculate p-values for RNA and protein chromosomal summary statistics in each sample, then combined using the Z-transformed method. Samples with a conservative combined p ≤ 0.0001 were annotated as aneuploid.

#### Virtual Karyotyping of Diversity Outbred Single-cell RNA-seq

Identification of aneuploid cells in single-cell RNA-seq data was performed using the R/CONICSmat package^63^. Each cell line was analyzed separately from the other 11 to keep expression comparisons limited to isogenic backgrounds. Cells with p ≤ 0.1 were annotated as aneuploid for Chromosomes 1, 6, 8, and 11.

#### Estimation of Aneuploid Cell Content in Bulk Data Sets

The fold-change in gene expression across duplicated chromosomes in aneuploid cells were quantified using the ‘detectBreakpoints’ function in the R/CONICSmat package. Cells with only one trisomic chromosomes were compared to cells with a completely euploid karyotype. A summary statistic was calculated by taking the median of all genes across the chromosome. The aneuploid content of bulk RNA-seq data can then be determined by the formula:

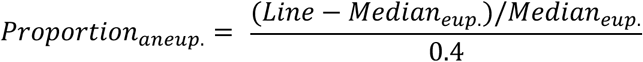

Where ‘*Proportion _aneup._*’ is the proportion of cells in the sample that have the chromosome in question duplicated, ‘*Line*’ is the summary statistic for the cell line whose aneuploid proportion is being calculated, ‘*Median _eup._*’ is the median of the summary statistic for lines annotated as euploid, and 0.4 is the proportional increase in expression expected if all cells in the sample were aneuploid for the chromosome in question, determined empirically.

#### Synteny Between Mouse and Human PSC Duplications

Identification of syntenic genes between mouse and human genomes was collected from The Jackson Lab Synteny Browser (syntenybrowser.jax.org). Visualization of connections between shared genes on chromosomes/segments duplicated in mouse and human PSCs was generated using the R/BioCircos package^64^. Mouse Chromosomes 1, 6, 8, and 11 were identified by this study. Human Chromosomes 1, 1q32, 7, iso7p, 12, 12p13.3, 14, 17, 17q25, 20, and X were identified from^1,3–8,65^.

#### Differential Expression Analysis

Differential expression was performed using R/Seurat package for single-cell RNA-seq. The function ‘FindMarkers’ was used with the default Wilcoxon rank sum test parameter to find differentially expressed genes between each comparison. GO term enrichment was performed using the R/ClusterProfiler package. Biological process ontologies of differentially expressed genes with log2 fold-change > |0.3| between trisomy 8 and completely euploid cells were visualized using the ‘gseGO’ function in the R/clusterProfiler package (minGSSize = 10, maxGSSize = 800, pvalueCutoff = 0.1) showing 16 categories split between activated and suppressed.

#### Genetic Mapping of Aneuploidy

Genetic mapping of aneuploidy states of the samples in the bulk data sets were performed using the R/qtl2 package^66^ using ploidy as a binary trait. Significance thresholds were calculated by running permutation analysis 1,000 times, generating a LOD = 8.74 for p = 0.01, LOD = 7.75 for p = 0.05, and LOD = 7.32 for p = 0.01.

#### X-Linked Gene Analyses

Correlations between transcript and protein abundance were calculated using the R/stats package. The 6 genes implicated in the sex differences in pluripotency of mESC were collected from^23^, the tumor suppressor genes on Chromosome X were collected from The Tumor Suppressor Gene Database (bioinfo.uth.edu/TSGene).

#### 2CLC Gene Expression in Bulk Data Set

Lists of genes upregulated in Zscan4^+^ cells in mESC cultures were collected from^39,40^ (supplemental data). Gene lists were filtered for genes with expression data in our bulk RNA-seq data set before calculating the expression z-scores for each remaining gene present across all samples. Significance between sum of z-scores for each chromosomal sex were calculated by t-test using the R/stats package.

#### LMO Demultiplexing and Preprocessing of Isogenic scRNA-seq

Data processing was performed mostly as described in the standard R/Seurat package tutorial. Only cells with less than 4% of total reads from mitochondrial genes, greater than 4,500 detected genes in the early data set and 4,000 detected genes in the aged data set were kept for further analysis. LMO barcode demultiplexing was performed using the ‘MULTIseqDemux’ function in the R/Seurat package. These cell identifications were found not to coincide with unsupervised clustering or XX biased gene expression in downstream analyses and were regarded as uninformative due to the low LMO barcode read counts.

#### Cell Cycle Scoring of Isogenic scRNA-seq

Cell cycle scoring was performed the same as in the DO mESC scRNA-seq data set – however, to ensure no subtle differences between chromosomal sexes due to cell cycling were observed, the difference between G2M and S phase scores were regressed out during scaling of the data.

#### Identification and Characterization of Pluripotent State, Differentiation Primed, MEF, and 2CLC Clusters

Pluripotent clusters were identified using expression of several key genes associated with general, naïve, and primed pluripotency. Additionally, several genes differentially expressed in XX and XY mESC were chosen to aid in identifying chromosomal sex. Genes were selected using information from^23,67–71^. MEF were identified using the expression of Thy1 and by predominant G1 cell cycle phase. 2CLC were identified using expression of the two Zinc finger and SCAN domain containing 4 (Zscan4) genes expressed in preimplantation development: *Zscan4c* and *Zscan4d*. Clustering resolution was set just sensitive enough that the 2CLC cluster separated out, as it was the smallest cluster of interest.

#### Virtual Karyotyping of Isogenic scRNA-seq

Cells with chromosomal CNV were identified using the same R/CONICSmat pipeline used for the DO mESC scRNA-seq data except that cells were not processed with the ‘SCTransform’ function.

#### Visualization

Most plots were generated using the R/ggplot2 package. QQ plots were generated using the R/ggpubr package, UpSet plots were generated using the R/UpSetR package, points underneath boxplots were plotted using R/ggplot2 with the ‘geom_quasirandom’ function from the R/ggbeeswarm package.

### mESC Cell Culture

The mESC used to validate the 2CLC differences between cell lines of varying chromosomal sex were derived from a cross between C3H/HeSn and Paj/J mice (JR #001529), designed to yield XO and XXY in addition to XX and XY embryos due to non-disjunction of the sex chromosomes. Three independently derived cell lines from each XX, XO, and XY embryos were obtained from the lab of Laura G. Reinholdt and maintained independently. Cells were cultured on plates coated in gelatin and irradiated MEF feeders and maintained on ESM (DMEM, 15% FBS, 1% Penicillin/Streptomycin, 1% GlutaMAX-1, 1% MEM NEAA, 100mM Sodium Pyruvate, 55 mM BME, 10^7^ units LIF). Cultures were tested for the presence of mycoplasma at the beginning and termination of the experiment. Pluripotency was assessed by germline chimera competence and cell identity by DNA sequencing.

### Flow Cytometry

A single-cell suspension was collected from each line by dissociation with 1X Accutase. Cells were stained for viability using Ghost Dye 780 (Tonbo Biosciences 13-0865-T100) on ice for 20 minutes. Immediately after, samples were fixed with 4% PFA (Electron Microscopy Sciences 15710-S) on ice for 15 minutes, quenched in subsequent washes with 0.125 mM glycine pH 3.0. Fixed cells were permeabilized in 1X Perm/Wash buffer (BD Biosciences 611202) on ice for 15 minutes. Samples were incubated in 1:6000 MERVL-gag primary antibody (EpiGentek A-2801-050) and 1:300 OCT 3/4 primary antibody (BD Biosciences 611202) at 4C for 1 hour. Samples were incubated in donkey anti-rabbit IgG Alexa 488 (Invitrogen A21206) and donkey anti-mouse IgG H+L Alexa 647 (Invitrogen A32787) secondary antibodies on ice for 20 minutes, then data was collected on the BD FAC Symphony A5.

## AUTHOR CONTRIBUTIONS

Diversity outbred mESC line derivation, culture, and collection: M.P., LGR, T.C. C3H/HeSn-Paf/J mESC line derivation: A.C, L.G.R. Mass spectrometry proteomics: S.P.G. RNA-seq and proteomics data set preprocessing: D.A.S., S.A. Single-cell RNA-seq data set: T.C. Computational analyses: A.S. with input from D.A.S., S.C.M. Human PSC karyotyping: K.L., S.T., E.M., T.L. Human PSC karyotype analysis: D.S. with statistical analysis by A.S. Manuscript writing and editing: A.S. and M.F.P. Study conception and design by M.F.P. and A.S.

## ACKNOWLEDGEMENTS

We would like to thank Gregory Carter and Christopher Baker of The Jackson Lab and Donna Slonim of Tufts University School of Medicine for consistent feedback throughout the investigation. We would like to thank Peter Andrews of Sheffield University for insights about differentially expressed gene sets in PSCs. Funding sources: The Jackson Laboratory.

## DATA AVAILABILITY

Accession numbers for RNA-seq and scRNA-seq data for these experiments are GSE287717 and GSE287621 respectively.

---

**Supplemental Figure 1.**
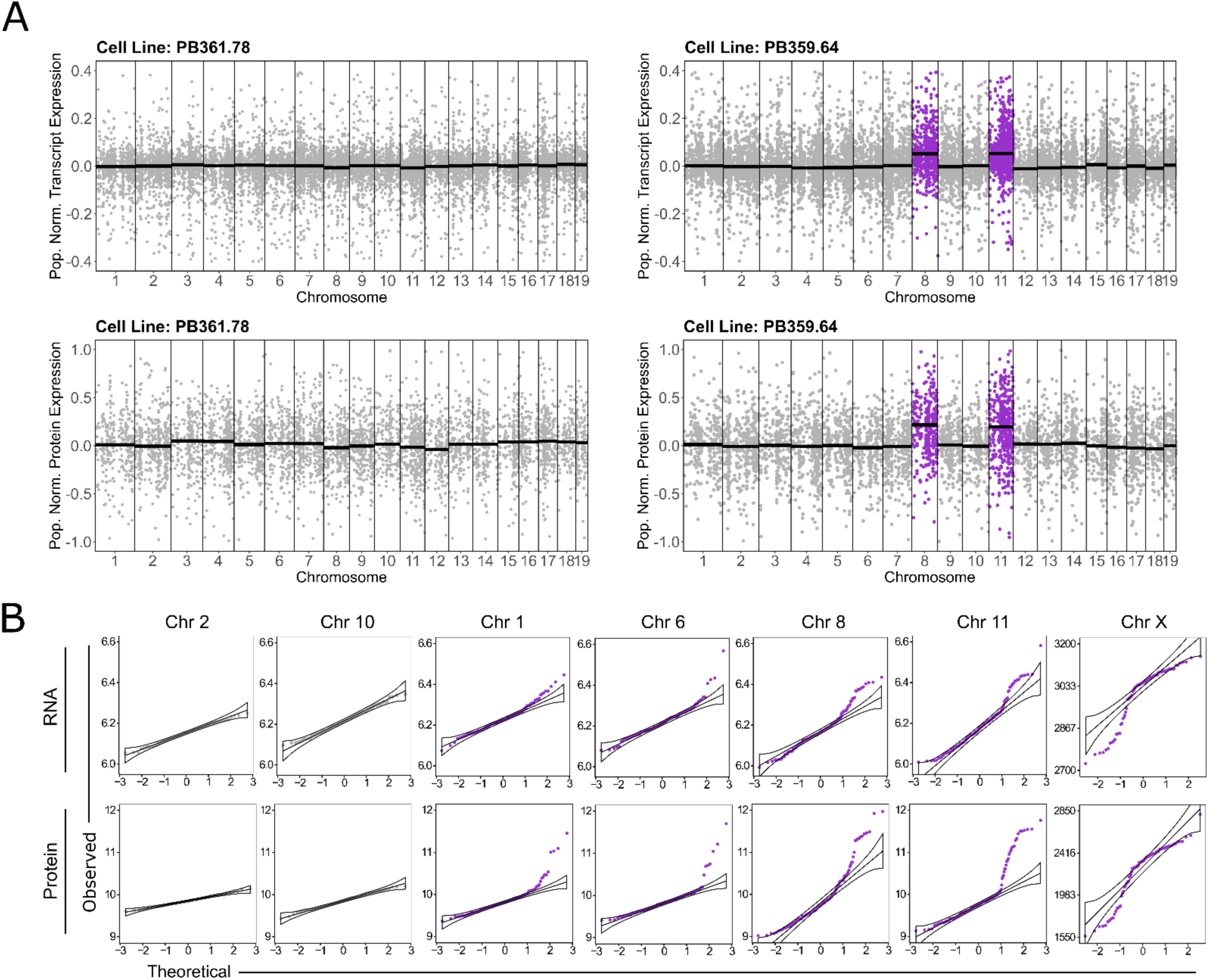
– Detecting aneuploid chromosomes in DO bulk RNA-seq and proteomics using consequent changes in gene expression. (A) Transcript and protein expression for each gene normalized to the population median for that gene in all mESC lines in the bulk data. Genes are plotted with equal spacing in order of their location on the chromosome. PB361.78 represents a cell line with no identified aneuploid autosomes, whereas PB359.64 represents a cell line with identified duplications for chromosome 8 and 11. Genes on euploid chromosomes are gray, genes on duplicated chromosomes are purple, and the black bars are the median normalized gene expression for each chromosome. (B) QQ plots comparing the distribution of median chromosomal gene expression – black bars in (A) – for the whole population to a theoretical normal distribution. Plots in gray are from commonly euploid chromosomes where no lines with copy number variations were found. Plots in purple are from chromosomes with non-normal distributions for transcript and protein, indicating lines had duplications or losses of that chromosome. For Chromosome X, only cell lines that were originally XX were compared.

**Supplemental Figure 2.**
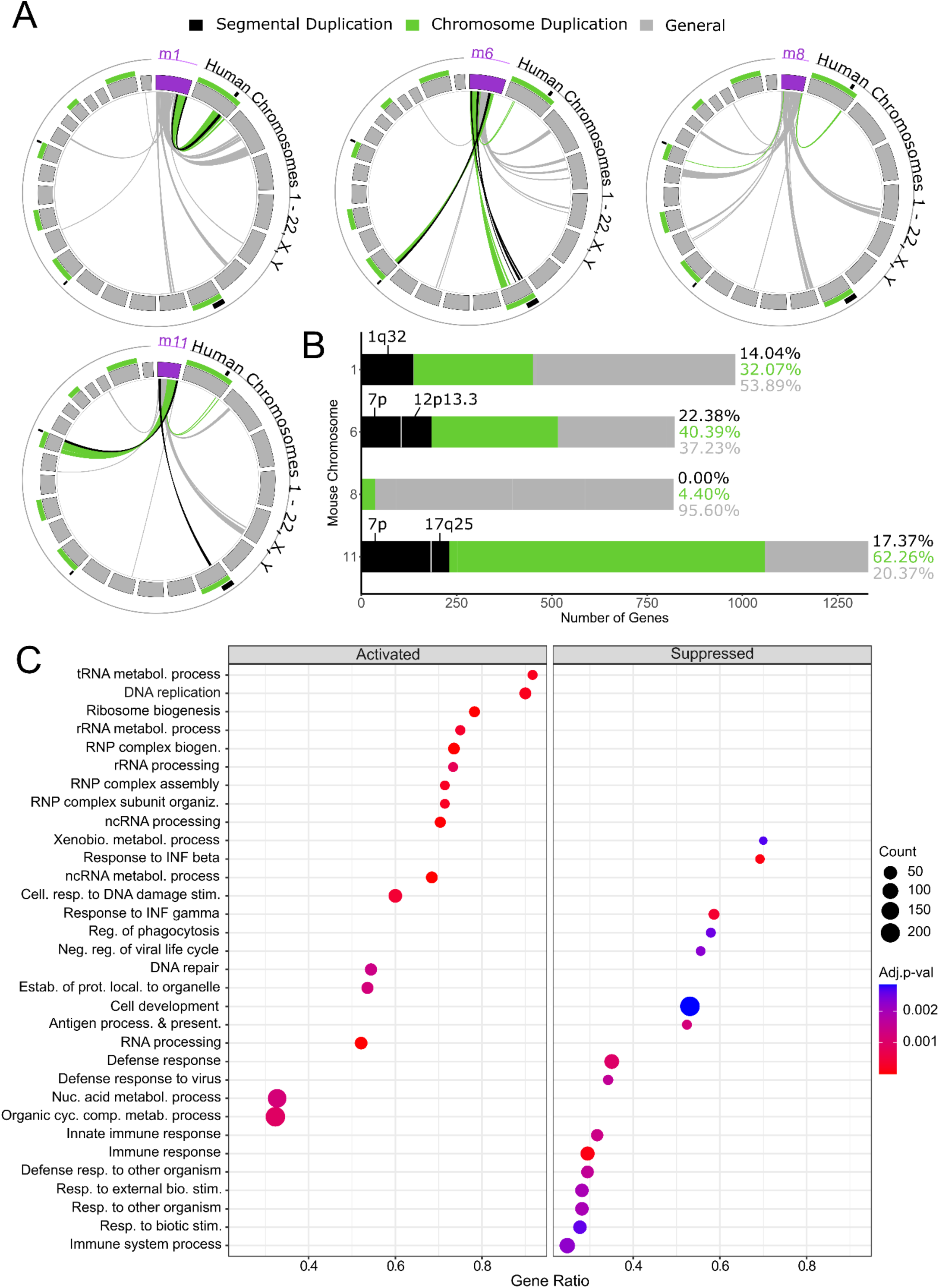
– Relating duplications identified in DO mESCs to commonly identified duplications in hPSCs. (A) Linking syntenic genes from their location on the duplicated mouse chromosome to their location in the human genome (syntenybrowser.jax.org). The duplicated mouse chromosome is represented in purple. Segments around the human genome and the links are colored by the type of duplication in hPSCs (black represents segmental duplications, green represents whole chromosome duplications, and gray represents general synteny – meaning no corresponding duplication). (B) The number of genes falling into each synteny category from each duplicated mESC chromosome. (C) Enriched GO terms for genes differentially expressed in cells with Chromosome 8 duplications compared to cells with normal karyotype in DO scRNA-seq.

**Supplemental Figure 3.**
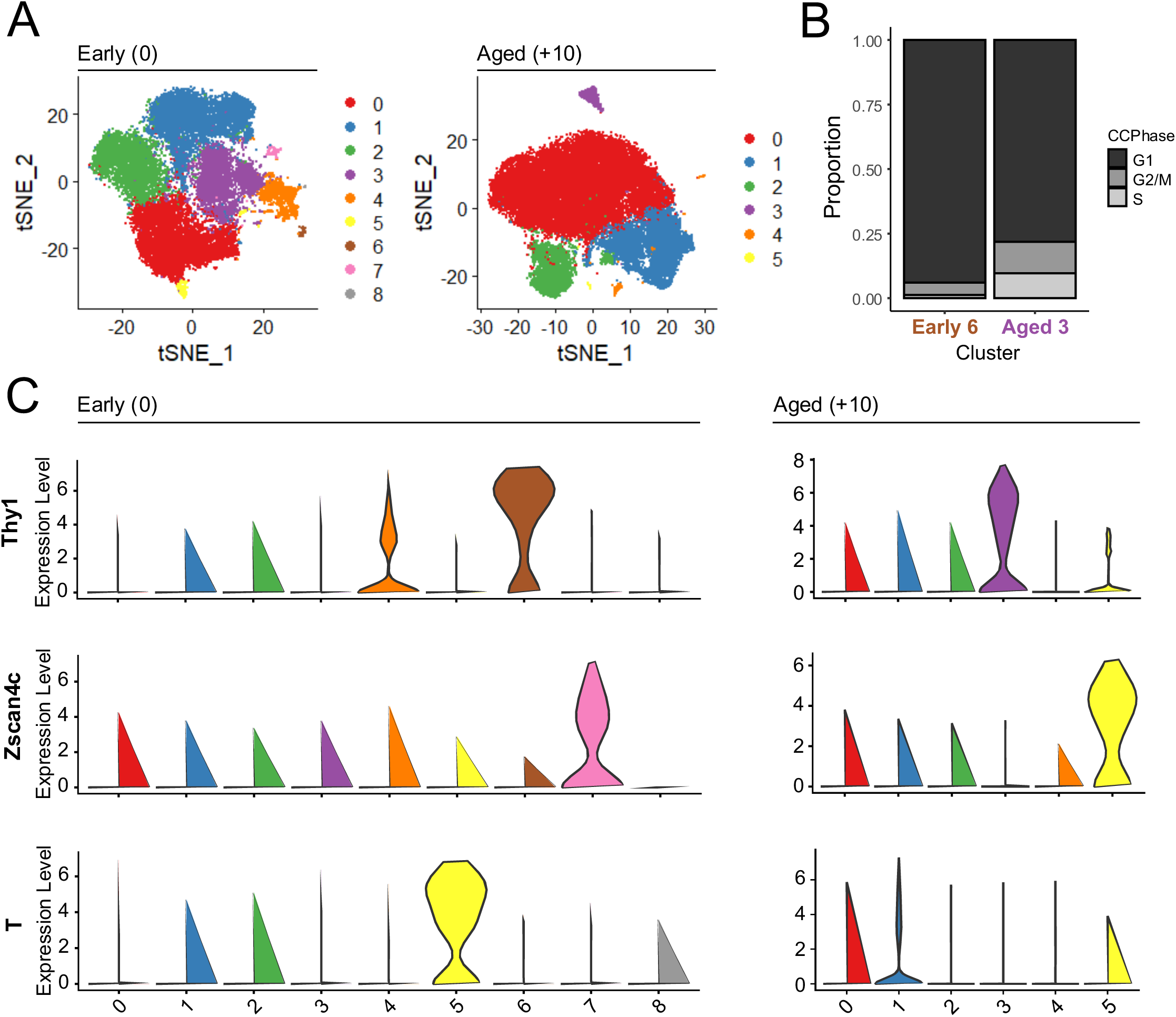
– Identifying MEF feeder, 2C-like state cell, and early differentiation clusters in scRNA-seq from isogenic cell lines by gene signatures. (A) Clustering of cells at first collection (early) and ten passages later (aged). (B) Predicted cell cycle phases of early cluster 6 and aged cluster 3 where cells were predominantly in G1, nonreplicating cells. (C) Cluster-level expression of genes marking MEF feeders (Thy1), 2C-like state cells (Zscan4c), and early differentiation (T).

**Supplemental Figure 4.**
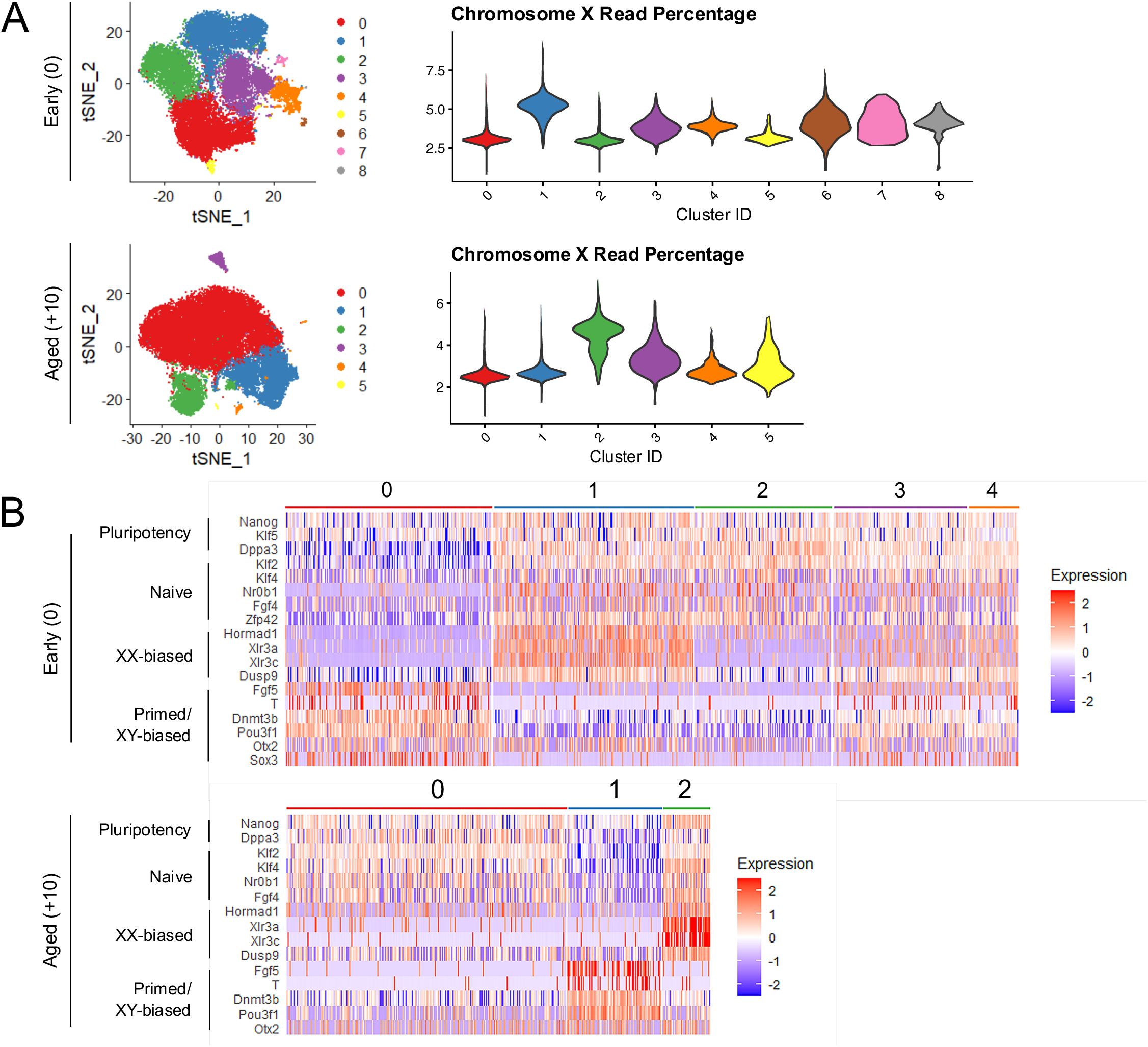
– Identifying XX, XO, and XY clusters in scRNA-seq from isogenic cell lines by gene signatures. (A) Percentage of total reads from Chromosome X in each cluster to identify the cells with one or two active X Chromosomes. (B) Relative expression of general and naïve pluripotency genes, genes with higher expression in XX mESCs, and primed pluripotency genes/genes with higher expression in XY mESCs. Only the remaining clusters not identified as MEF feeders, 2C-like state cells, and non-pluripotent cells were compared.

